# Cortical neural activity predicts sensory acuity under optogenetic manipulation

**DOI:** 10.1101/119453

**Authors:** John J. Briguglio, Mark Aizenberg, Vijay Balasubramanian, Maria N. Geffen

## Abstract

Excitatory and inhibitory neurons in the mammalian sensory cortex form interconnected circuits that control cortical stimulus selectivity and sensory acuity. Theoretical studies have predicted that suppression of inhibition in such excitatory-inhibitory networks can lead to either an increase or, paradoxically, a decrease in excitatory neuronal firing, with consequent effects on stimulus selectivity. We tested whether modulation of inhibition or excitation in the auditory cortex could evoke such a variety of effects in tone-evoked responses and in behavioral frequency discrimination acuity. We found that, indeed, the effects of optogenetic manipulation on stimulus selectivity and behavior varied in both magnitude and sign across subjects, possibly reflecting differences in circuitry or expression of optogenetic factors. Changes in neural population responses consistently predicted behavioral changes for individuals separately, including improvement and impairment in acuity. This correlation between cortical and behavioral change demonstrates that, despite complex and varied effects these manipulations can have on neuronal dynamics, the resulting changes in cortical activity account for accompanying changes in behavioral acuity.

**Author summary:** Excitatory and inhibitory interactions determine stimulus specificity and tuning in sensory cortex, thereby controlling perceptual discrimination acuity. Modeling of such excitatory-inhibitory circuits has predicted that suppressing the activity of inhibitory neurons can lead to increases or, paradoxically, decreases in excitatory activity, depending on the architecture and modulation parameters of the inhibitory component of the network. Here, we capitalized on differences between subjects to test whether suppressing/activating inhibition and excitation across a range of parameters in sensory cortex can in fact exhibit such paradoxical effects for both stimulus sensitivity and behavioral discriminability. Indeed, we found that the same optogenetic manipulation in the auditory cortices of different mice could improve or impair frequency discrimination acuity, in a fashion that was predictable from the effects on cortical responses to tones. The same manipulations sometimes produced opposite changes in the behavior of different individuals, supporting theoretical predictions for inhibition-stabilized networks.

## 1 Introduction

Sensitivity to sensory signals depends on neuronal tuning to specific parameters of sensory stimuli, such as orientation of edges for visual stimuli, or tone frequency for auditory stimuli. Such neuronal selectivity arises in many brain areas and is shaped by complex, interconnected circuits of excitatory and inhibitory neurons [1]. The balance between inhibitory and excitatory stimulus representation in the sensory cortex has been proposed to underlie learning- and adaptation-dependent changes in stimulus-driven responses [2]. Recently, a number of studies have begun to unravel the role of inhibition in sensory processing, empowered by recently developed optogenetic techniques [for example 3, 4-10]. These methods drive specific classes of neurons to express an opsin, such that shining light over that brain area either activates or suppresses neuronal activity selectively [11-15].

While optogenetic techniques provide exquisite molecular and temporal specificity for testing the function of specific cell types in sensory acuity, they currently carry inherent and technical limitations that can lead to different levels of opsin expression and activation or suppression strength across individual animals. Since measurements are typically performed in multiple animals, these specific differences are accounted for using statistical analyses while summarizing the average effects of optogenetic perturbations. We postulated that these differences could be instead *exploited* in order to characterize the diversity of effects across subjects, thereby deepening our understanding of the function of inhibitory-excitatory interactions in sensory processing.

Theoretically, it has been predicted that within balanced excitatory-inhibitory circuits, increasing inhibition can either decrease excitatory neuronal activity or, paradoxically, increase it, depending on specific perturbation parameters and circuit properties [16]. We therefore hypothesized that the differences in technical parameters of optogenetic stimulations across animals could evoke both positive and negative effects on the firing rate of the target neuronal population. Focusing on tone frequency representation in the auditory cortex [9], we tested whether this was indeed the case by up- or down-regulating the activity of either inhibitory or excitatory neurons and measuring the resulting changes in frequency discriminability based on neuronal population activity. Our previous analyses had demonstrated that, on average, suppressing the activity of the most common class of interneuron, parvalbumin-positive neurons (PVs), in auditory cortex (AC) impaired behavioral frequency discrimination acuity, while activating PVs improved it, whereas activating excitatory neurons did not have an effect [9]. Here, we compared changes in neuronally predicted and behavioral frequency discrimination acuity due to optogenetic manipulations.

Optogenetic interventions led to changes in tone-evoked responses of recorded neurons. As predicted, the same manipulation sometimes produced opposite changes in neurometric sensitivity and in behavior for different individuals. Computational analysis predicted consequent changes in discrimination thresholds, which explained the measured behavioral changes, including improvement or impairment of discrimination driven by the same optogenetic manipulation in different individuals. Our results thus demonstrate that, although these manipulations have complex effects on the neural network, the resultant changes in activity are sufficient to predict changes in behavior.

## Results

### Measuring frequency discrimination acuity

To measure frequency discrimination acuity, we used a procedure based on pre-pulse inhibition of the acoustic startle reflex (Figure 1A, B). Like other mammals, mice startle to loud noise. The startle response, as measured by the change in pressure exerted by the subject on a balance platform, is typically decreased when the startle noise is preceded by a brief tone that mice can detect – a phenomenon termed pre-pulse inhibition (PPI). We presented mice with a continuous tone at one frequency, which was stepped to a tone of a different frequency just prior to the startle noise. The startle response was attenuated as the frequency difference between background and pre-pulse tones increased (Figure 1B, C). This attenuation reflects the ability of the mouse to detect frequency differences [9, 17]. We characterized frequency discrimination acuity in terms of a *behavioral threshold* -- the frequency difference that produced 50% of the maximum PPI (Figure 1 C).

**Figure 1:**
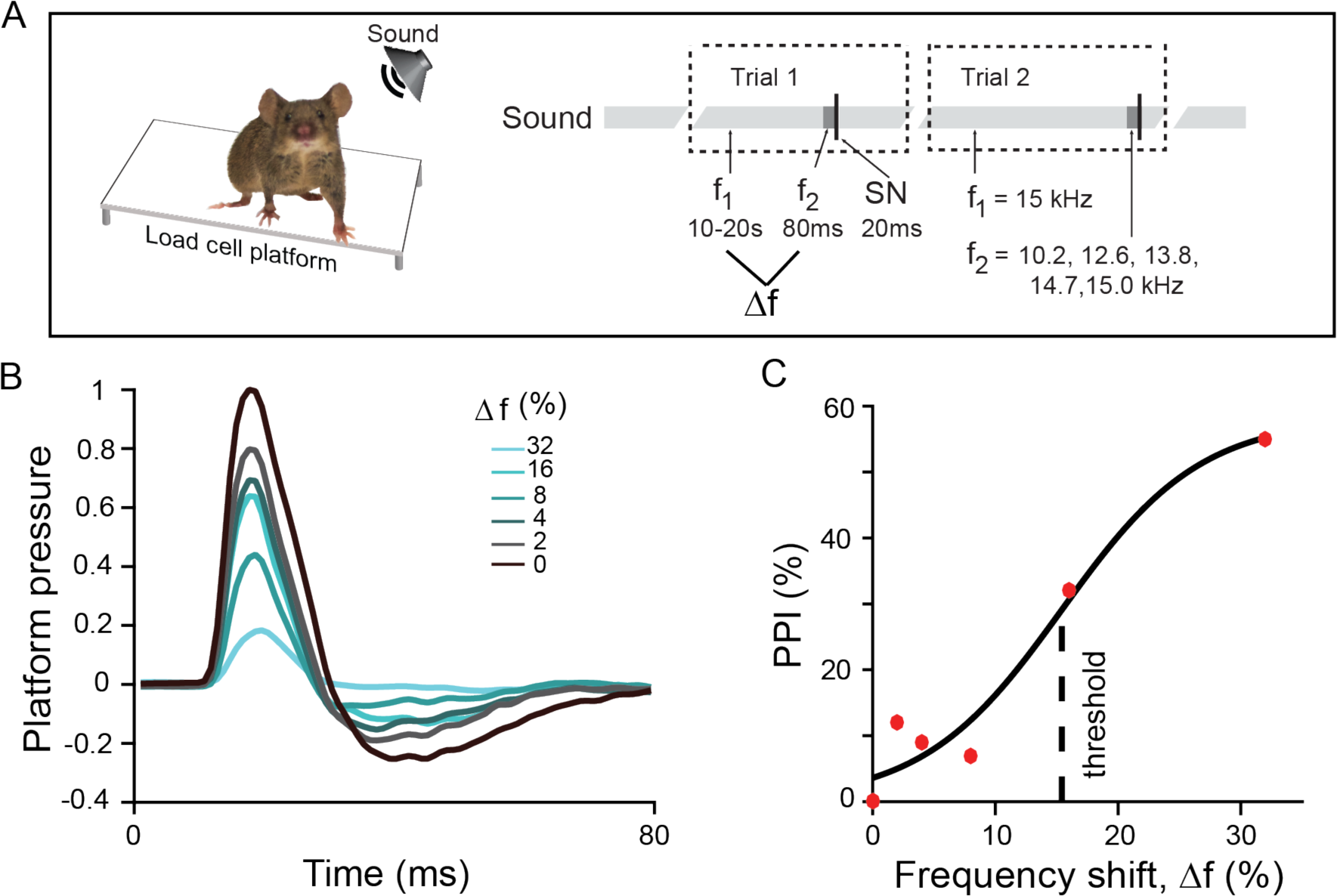
Measurement of behavioral frequency discrimination acuity. **A**. Schematic of measurement of frequency discrimination acuity in mouse. **Left**: Startle response is measured as pressure the subject exerts on a platform. **Right**: Sound stimulus time course: an ongoing background tone (light gray, f_1_) is followed by a brief pre-pulse tone of different frequency (dark grey band, f_2,_) and then by a startle noise (thin black band, SN). **B**. Normalized time course of platform pressure during the startle response to noise for different pre-pulse tones for an exemplar mouse. Time relative to SN onset. **C**. Pre-pulse inhibition measured as reduction in the acoustic startle response as a function of the frequency shift (Δf) between the background and pre-pulse tones (see Methods for definition) of an exemplar mouse. PPI does not reach 100% because even with an easily identifiable prepulse tones, the animal still startles. Dots: data, solid line: fit.

### Neurometric discrimination thresholds from Fisher Information

Next, we recorded the activity of putative excitatory cells (Methods) in AC, while presenting the awake head-fixed mouse with a random tone pip stimulus (50 ms tone pips presented every 500 ms, frequency changed at random, Figure 2A). For each frequency-tuned neuron, we measured frequency response curves (mean firing rate as a function of tone frequency, Methods and Figure 2B). To estimate a discrimination threshold from the frequency-tuned neural population recorded in each mouse we used Fisher Information analysis. To do so, we fit Gaussian functions to the response curve of each neuron, and assumed that neurons responded independently and with Poisson variability. Standard methods then provided the Fisher information for frequency discrimination based on the population response (Methods; Figure 2C) [18]. Fisher information quantifies the amount of information that neural responses provide to distinguish nearby frequencies. Decoding sensitivity increases with Fisher information. Since this information is large when the neural response changes quickly as a function of frequency, it is higher on the slopes of the tuning curves than in the center.

**Figure 2:**
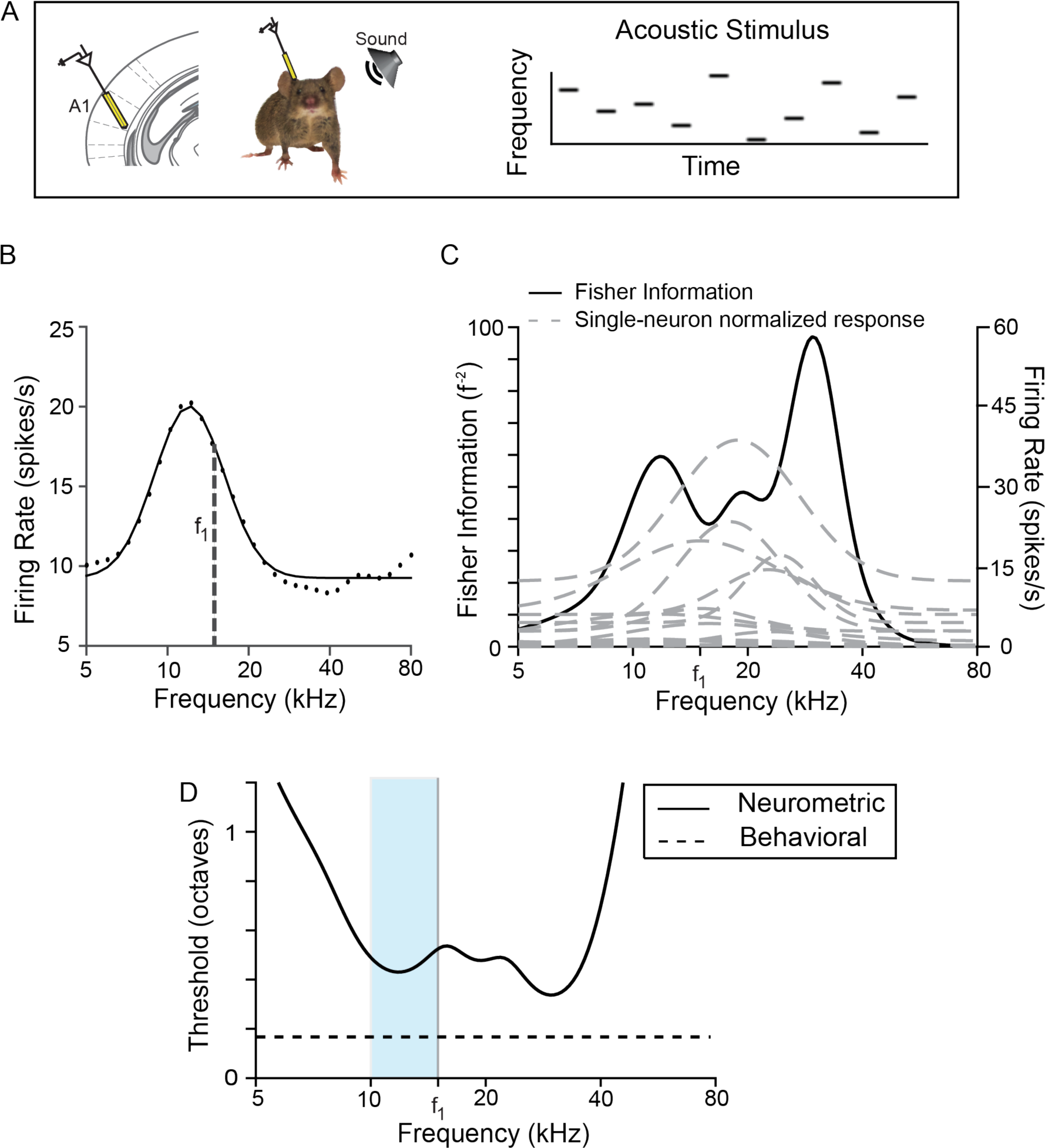
Measurement of neurometric frequency discrimination acuity. **A**. **Left**: Schematic of electrophysiological recording of neuronal responses in the primary auditory cortex (A1) in awake mouse. **Right**: The stimulus consists of a pseudorandom sequence of pure tones at varying frequency and intensity levels. **B**. Representative frequency response function for a single neuron (f_1_ = background tone in Fig. 1). Black dots: data, black line: fit. **C**. Fisher information for tone discrimination (black) computed on the basis of frequency response functions (gray dashed) of neurons (N=14) recorded in the same mouse. **D**. Neurometric threshold for decoding frequency (solid) computed on the basis of the inverse square root of Fisher Information computed in **C**. The neurometric threshold based on the recorded population lies above behavioral threshold for discrimination around f_1_ (dashed line). Light blue band indicates the region in frequency space from which behavioral measurements were taken.

The inverse square root of the Fisher Information bounds the accuracy with which nearby frequencies can be distinguished based on population activity. This quantity is by definition the *neurometric threshold*, and gives the frequency difference that can be discriminated with 70% accuracy. Figure 2D shows the neurometric thresholds determined in an individual mouse on the basis of the neural population recorded in its cortex.

A direct comparison of the neurometric threshold with behavior faces three challenges: (a) discrimination accuracy need not translate linearly into thresholds determined from the pre-pulse inhibition of the acoustic startle response measured here, (b) cortical recordings inevitably sub-sample the population of responsive neurons, and (c) frequency discrimination is supported by multiple pathways, some of which may not involve the auditory cortex. In general, we expect the neurometric thresholds computed here to be higher than the behavioral thresholds, because of the limited number of recorded neurons (e.g., Figure 2D).

In view of these challenges we did not seek to predict absolute behavioral thresholds from the population of recorded frequency-tuned neurons. Rather, we made a differential estimate: we manipulated excitatory and inhibitory neuronal activity in the auditory cortex optogenetically, and tested whether there was a correlation between the resulting *changes* in the neurometric estimate of discrimination thresholds and corresponding changes in behavior.

### Optogenetically induced changes in neurometric and behavioral thresholds are correlated

Frequency tuning of AC neurons is thought to depend on the combination of excitatory and inhibitory inputs [9, 19-22]. To manipulate tone responses of AC neurons, we targeted the most common interneuron subtype, parvalbumin-positive interneurons (PVs). We drove PVs to express Archaerhodopsin (Arch) or Channelrhodopsin (ChR2) by injecting a floxed Arch- or ChR2-encoding virus in PV-Cre mice. We verified the efficiency of the viral transfection and light stimulation by measuring the effect of light on spontaneous firing rates of neurons. As expected, when PVs were activated by light, thus increasing inhibition, spontaneous activity decreased in the recorded population (Figure 3A) [10]. Optogenetically suppressing PVs had the opposite effect: the spontaneous rate of most recorded neurons increased (Figure 3B). We also drove excitatory neurons in AC to express ChR2 and found that optogenetic illumination of AC led to an elevated spontaneous firing rate (Figure 3C). Thus, we reliably used light to manipulate the activity of excitatory and PV neurons in AC.

**Figure 3:**
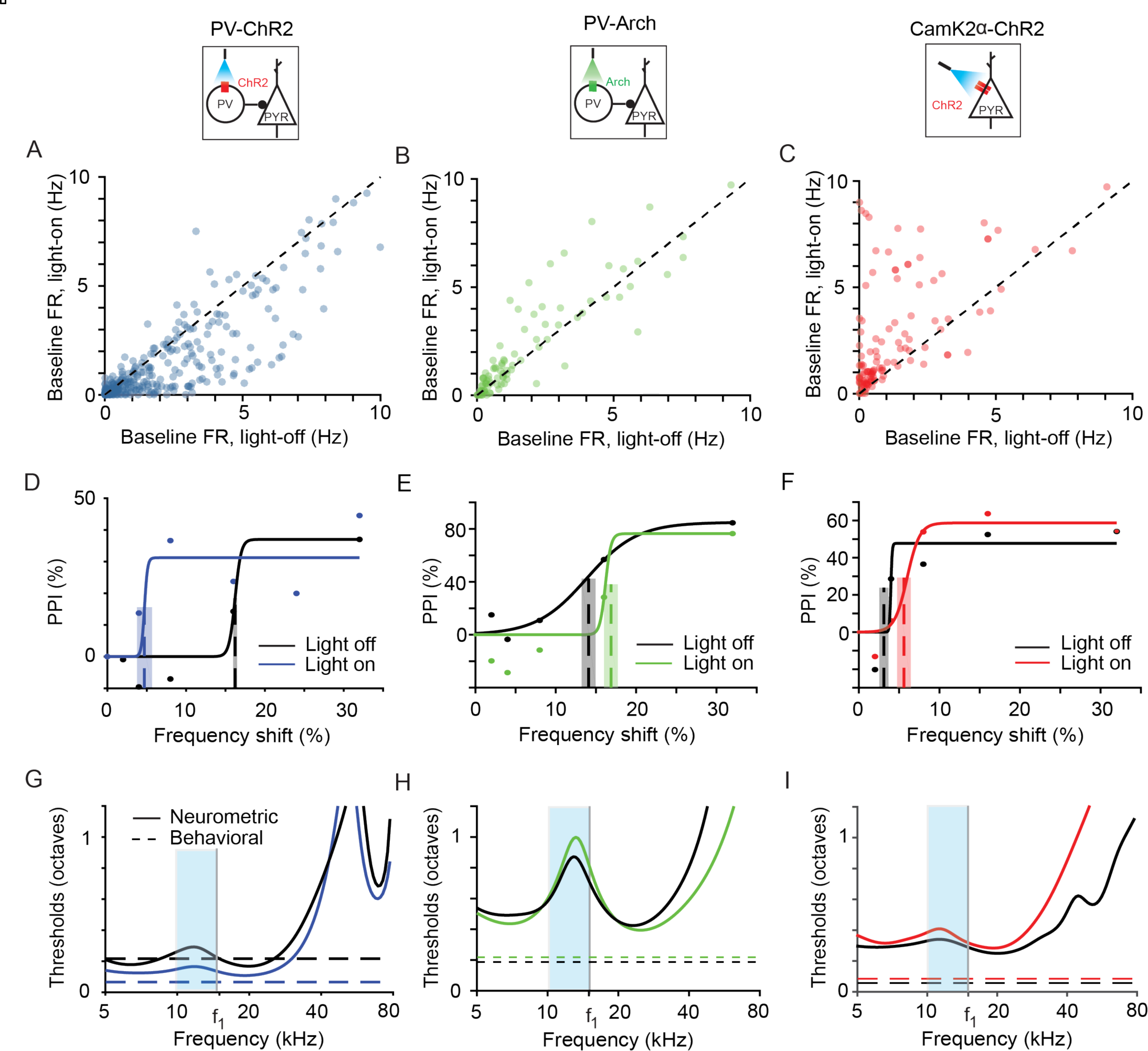
Optogenetic manipulation of PV activity shifts behavioral and neurometric frequency discrimination thresholds in individual subjects. **A,B,C**: Baseline firing rate of light-on versus light-off trials for all frequency-tuned neurons pooled across subjects in PV-ChR2 (Blue), PV-Arch (Green), CamK2a-ChR2 (Red) mice, respectively. **D,E,F**: PPI as a function of tone frequency shift for exemplar mice. Best estimated thresholds (dashed lines) and uncertainties (overlaid gray rectangle) are plotted for reference. Black: light-off trials; Blue, Green, Red: light-on trials. Dots: data, solid lines: best fit curve. **G,H,I**: Neurometric threshold estimate as inverse square root of Fisher information (solid) and behavioral threshold at *f*_1_, (horizontal dotted) for the same mice as **D,E,F**. Light blue bands indicate the region in frequency space from which behavioral measurements were taken.

We next measured the behavioral effects of manipulating neural activity in AC. On half of the trials, we illuminated AC at the same time as providing the pre-pulse stimulus. Activating PVs (Figure 3D) decreased the threshold for most animals (N=5) and increased it for some (N=2), while a few (N=4) did not have a statistically significant threshold change. Suppressing PVs (Figure 3E) produced an increase in the frequency discrimination threshold for two animals, and did not produce statistically significant change for one animal. Optogenetically activating excitatory neurons increased the threshold for most animals (N=3) and decreased it for one, with one animal displaying no threshold change (Figure 3F). For several animals, (N=6) the manipulations did not produce any significant behavioral changes.

Finally, we measured the effect of manipulating neuronal activity on the neurometric frequency discrimination thresholds predicted from the recorded population. Activating PVs led to a decrease in the predicted threshold (Figures 3G, 4A, B, blue) for most PV-ChR2 mice (N=7) and an increase for some (N=2). The predicted threshold increased for three PV-Arch mice after suppressing PVs (Figures 3H, 4A, B green), but decreased for two. Activating excitatory neurons (Figures 3I, 4A, B, red) increased the predicted threshold for all CamKIIa-ChR2 mice (N=3). There was no predicted change for two mice (<2% change in predicted threshold).

A potential confound for optogenetic manipulation of cortical activity is the scattering of light through the tip of the optic fiber. To prevent the light manipulation to serve as an additional prepulse, we placed a bright LED in the chamber, which served to adapt the retina of the mouse to scattered light from the optic fiber [23]. Another limitation of light-driven manipulations of neuronal activity is a potential cascade of prolonged circuit-level dynamics. Therefore, to facilitate the comparison of the effect on light-off and light-on trials, light was presented on light-off trials as well, but not concurrent with the prepulse and startle stimuli. Indeed, light presentation alone did not affect the startle response (PV-ChR2 mice, blue light: N = 11, p>0.05; PV-Arch mice, green light: N = 8, p>0.05) and thus did not serve as a pre-pulse of the acoustic startle response. As an additional control, we injected a cohort of mice with virus carrying only the fluorescent reporter, and not the opsin. In these animals, there were no significant effects on frequency discrimination acuity (N = 6, p>0.05) [9]. These two aspects of stimulus design and control measurements thus confirmed that the observed behavioral changes were unlikely due to artifacts of light presentation, but rather due to light-driven manipulation of cortical activity.

For most individuals (N=15/19) and on average, the sign of the neurometric change in threshold with the optogenetic manipulations matched the sign of the change in behavioral threshold (Figure 4A, B). This qualitative agreement was striking, given that the electrophysiological recordings only sample a few neurons, while the light has a global effect on the auditory cortex and sometimes leads to opposite behavioral changes. To quantify the correlation, we compared the neurometric and behavioral frequency discrimination thresholds for each mouse, under light-on and light-off conditions. The number of recorded tone-responsive neurons varied significantly between mice (14-104 per animal). Using the linear dependence of the Fisher Information on population size we estimated that ∼1000 independent neurons would be necessary for the neurometric thresholds to match the absolute behavioral thresholds (see Methods). This number differed between different mice, presumably because of the limited sampling, although was consistent in order of magnitude. Since optimizing the size of the population for each mouse would have skewed analysis across subjects, we divided the Fisher Information computed from the population by the number of neurons, and then scaled the result back linearly to the same effective population size for all mice (N = 1000 units).

**Figure 4.**
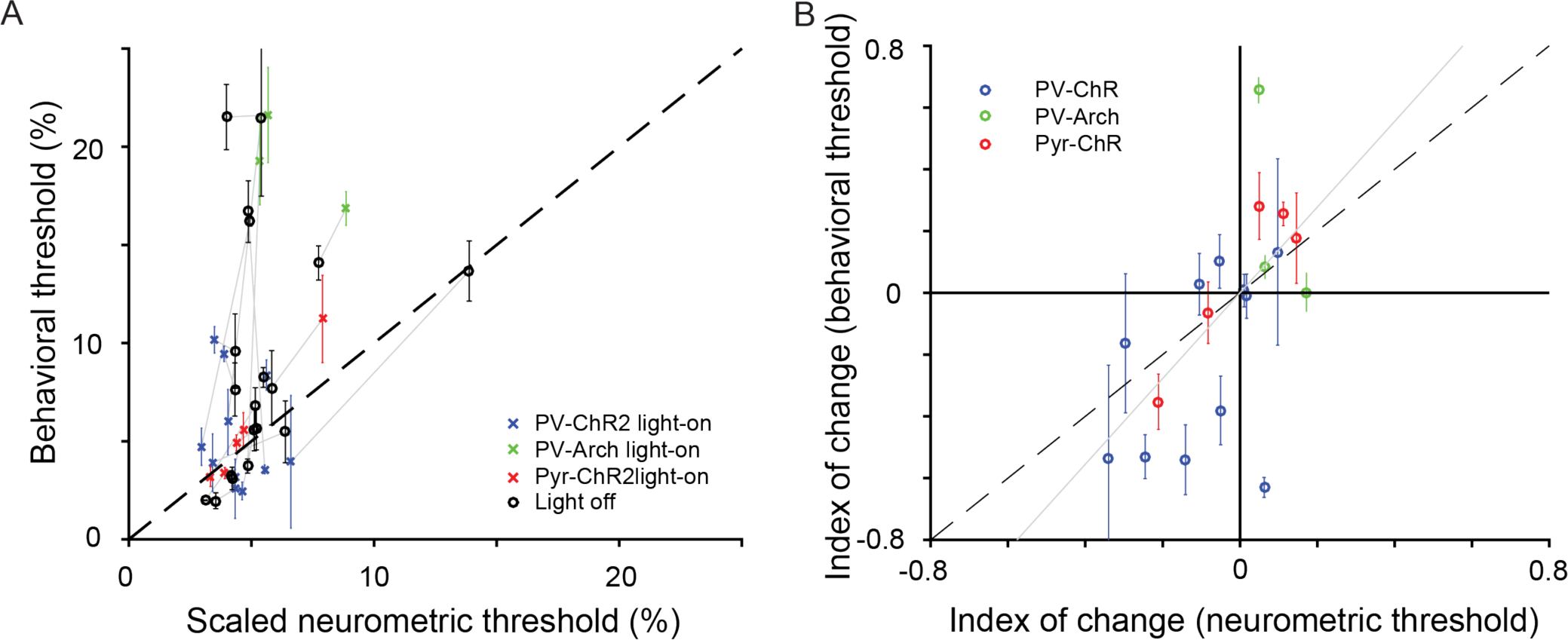
Changes in A1 tone responses due to optogenetic manipulations predict changes in behavioral frequency discrimination acuity across individuals. **A**. Behavioral versus scaled neurometric frequency discrimination thresholds. Neurometric threshold (computed as inverse of Fisher information squared for tone-evoked responses from all neurons recorded in each mouse) is scaled to an effective population size of 1000 neurons to control for differences in numbers of measured neurons. Changing this scale factor is equivalent to changing y-axis labels. The scaled neurometric threshold based on the small recorded population was significantly (but weakly so, C=.37, p=0.02) correlated with the behavioral threshold (computed as the shift in frequency between the background and pre-pulse tone that evoked 50% of the maximum PPI). Each of 19 mice contributes 2 data points, representing the threshold computed on the basis of light-on and light-off trials. Gray lines connect light-on and light-off estimates for each mouse. **B**. The index of change in neurometric threshold (difference between thresholds computed from data on light-on vs. light-off trials divided by the sum) was strongly correlated with the behavioral frequency discrimination (Correlation coefficient = 0.59, p=0.007). There is one data point for each mouse. Gray line is the best fit line through the origin. Behavioral errors were computed as described in the Methods.

We next compared the resulting estimate to the behavioral threshold. We first found a correlation between the absolute behavioral and neurometric thresholds under all conditions (Figure 4A, S2 Figure, S1 Table). The correlation was statistically significant (*C* = .37, *p* = .02, *N* = 38, including a light-on and a light-off measurement for each of 19 mice), but only weakly so. To test more closely the effect of the optogenetic manipulations, we computed an index of change as the difference in thresholds before and after application of light, divided by the sum. We found that index of change of the neurometric thresholds was strongly correlated with the behaviorally measured index of change in frequency discrimination acuity threshold (Figure 4B, *C* = .59, *p* = .007, *N* = 19).

These correlations suggest that: (a) auditory cortex does modulate frequency discrimination behavior, (b) the effects seen in the small recorded patch are representative, and (c) individual differences in auditory behavior are directly driven by differences in excitatory and inhibitory interactions in cortical circuits.

### Controls: effects of optogenetic manipulation on neural variability and correlations

Our model makes two assumptions that could be violated in neural systems: that cortical neurons obey Poisson statistics and that neural responses are independent of one another. In order to test the first assumption in our data, we measured the Fano factor of the recorded neurons. A large Fano factor indicates high neuronal variability [24]. The average Fano factor was around 1.2, similar to the value expected for Poisson neurons and to that previously measured across different cortical areas [25]. We found that none of the three optogenetic manipulations (PV-ChR2: *t*_335_ = .4, *p* = .69; PV-Arch: *t*_89_ = .92, *p* = .36; Pyr-ChR2: *t*_133_ = −.2, *p* = .84; Figure 5A-C) had a systematic effect on the distribution of Fano factors. This justified the original analysis, in which neurons were treated as Poisson, both with and without optogenetic manipulation.

**Figure 5.**
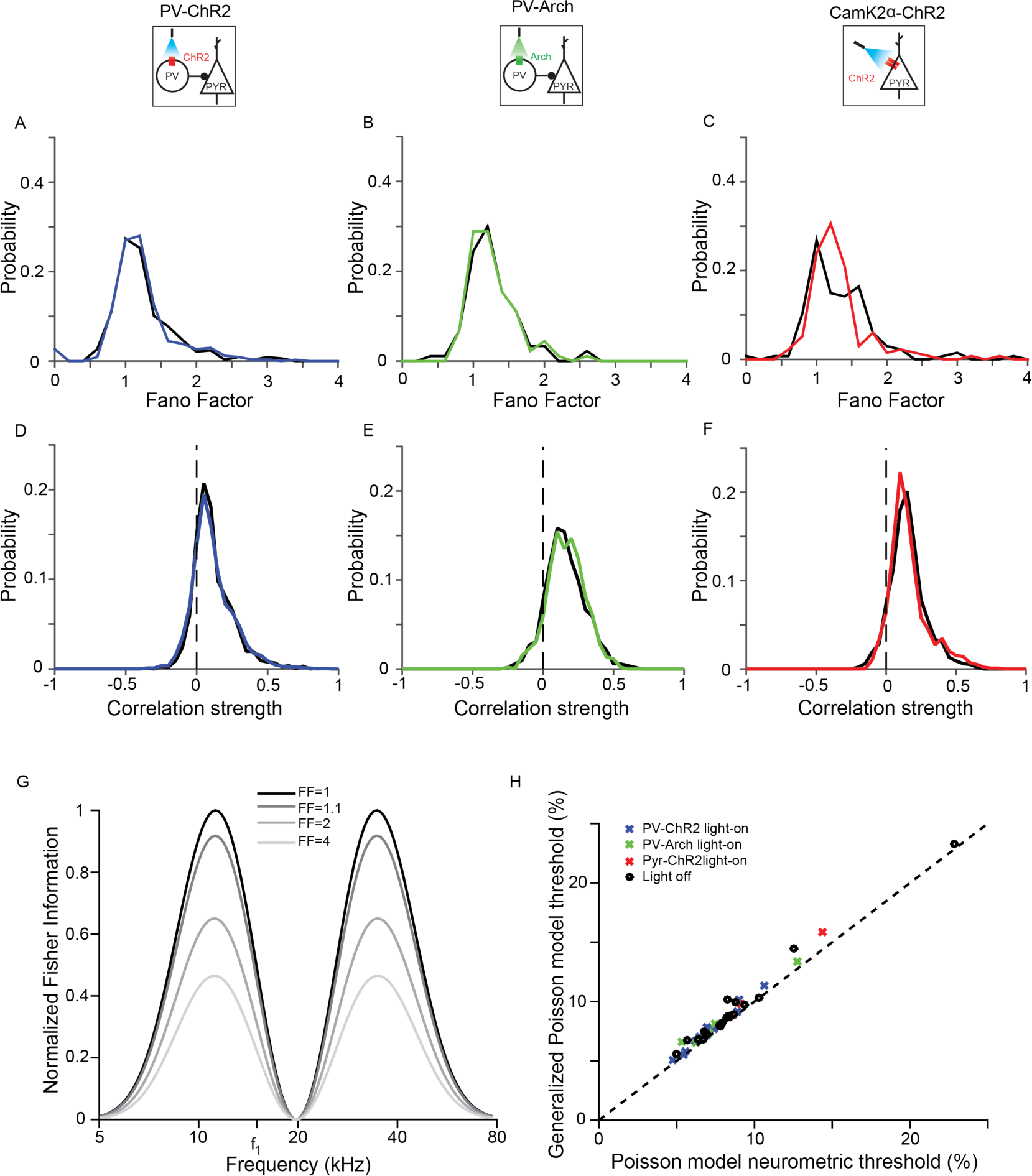
Optogenetic manipulations do not change neuronal variability or correlations. **A-C**. Fano factor pooled across mice distributions are similar under light-on and light-off conditions. A: PV-ChR2, B: PV-Arch, C: CamK2a-ChR2. Black: light-off trials; Blue, Green, Red: light-on trials. Dots **D-F**. Pairwise correlation distributions pooled across mice are similar under light-on and light-off conditions. D: PV-ChR2, E: PV-Arch, F: CamK2a-ChR2. Colors same as in A. **G**. Increasing Fano factor reduces Fisher Information, shown here for a single neuron with Gaussian tuning curve (amplitude = 8 spikes/s, center frequency 20kHZ, tuning width = 0.2 decades) with a constant baseline (2 spikes/s). **H**. Incorporating the measured Fano factors into our model of neuronal firing via a generalized Poisson model has a weak effect on the predicted threshold.

We next considered that neural variability has been observed to increase with the activity level of neurons [26]. It is plausible that the most active neurons, which have the largest impact on Fisher information, may be more variable than expected from the average Fano factor value. The Fisher information decreases as the Fano factor increases (Figure 5G), so higher variability in the most active neurons would disproportionately decrease the neurometric discriminability. To test whether this might be the case, we computed the neurometric thresholds using a generalized Poisson noise model that took into account the Fano factor of each recorded neuron separately. Neurometric thresholds using this model changed only slightly from the thresholds computed using the simple Poisson noise model (Figure 5H), again justifying our analysis.

We also measured the strength of pairwise neural correlations, and observed they tended to be small but positive (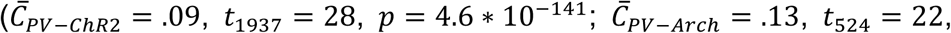 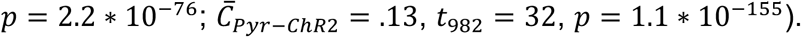 The optogenetic manipulations had no systematic effect on the distribution of correlations (paired *t* test ns: PV-ChR2 *t*_1937_ = .26, *p* = .80; PV-Arch *t*_524_ = −1.3, *p* = .18; Pyr-ChR2 *t*_982_ = −1.7, *p* = .09; see Figure 5D-F, S1). Correlations in similar models have been observed to lead only to small increases in population-level discrimination threshold [27]. The small changes in threshold observed in previous work, in addition to the negligible change in the correlation distribution that we observe upon optogenetic manipulation, make it unlikely that correlations account for the neurometric threshold changes that we see when cortex is manipulated.

## Discussion

Theory has shown that providing inputs to inhibitory neurons in a balanced excitatory-inhibitory network can lead to either a decrease, or, paradoxically, an increase of excitatory responses, depending on the specifics of recurrent coupling in the network and the strength of the manipulation of inhibitory activity [16]. By examining changes evoked by optogenetic manipulations across subjects, we find that modifications of the excitatory-inhibitory interactions in auditory cortex drive diverse, and sometimes opposite, changes in tone-evoked responses across individuals. Remarkably, we found a strong correlation between these changes in neuronal populations, and behavioral changes in the acuity of frequency discrimination by individual mice. Thus our results demonstrate that the same optogenetic manipulations of excitatory and inhibitory neuronal activity in different individuals, can have diverse effects on sensory discrimination. More generally, these findings support a role for excitatory-inhibitory networks in the cortex in mediating sensory discrimination.

Where does this variability in the effects of manipulations across animals arise from? Some of the differences may be due to the inherent variability in circuitry across animals: the excitatory-inhibitory circuit can have different connectivity patterns, and animals may exhibit differences in frequency sensitivity due to a combination of genetic and environmental factors. In addition, optogenetic manipulations introduce variability across animals due to technical limitations of the technique. In testing the function of specific neuronal cell types in sensory processing, there are a number of potential confounds. For example, the exact position of the needle for virus injection within AC, the depth of penetration, and the spread of the virus, can all skew the extent to which the virus is expressed within AC. The position of the optic cannula relative to the spread of the virus injection as well as the recording electrode would affect how many neurons are stimulated and how strongly, and also how many of those neurons are being picked up by the recording. Small changes in any of these parameters can potentially lead to strong differences in the functional effects of optogenetic perturbations, activating a different fraction of neurons across different laminae, and in different tonotopic regions of the cortex. In this study, we capitalized on these differences because they allowed us to assay the range of potential effects of optogenetic manipulations. It was critical, however, to ensure that within animals, optogenetic manipulation would produce the same effect during behavioral testing and during electrophysiological recordings. We therefore used an implanted optic cannula and a light delivery system with the same settings to deliver light both during the electrophysiological recording and the behavioral measurements, so that stimulation parameters were the same for both.

Our results provide insight into potential role of cortical inhibition in shaping frequency discrimination behavior. Cortical inhibitory neurons have been hypothesized to modulate numerous aspects of tone-evoked responses in the excitatory cortical circuit, such as tuning width, reliability of firing, tone-evoked response strength, and correlations in firing rate activity [2, 19, 28]. Neurons in the auditory cortex change their tuning properties with learning, attention or experience [29-35], suggesting that these changes can underlie changes in auditory perception. Indeed, AC was shown to play an important function in enabling learning-driven changes in auditory processing [17, 36]. Our present work demonstrates that modulation of excitatory-inhibitory dynamics can in principle support a wide range of modulatory effects on auditory frequency discrimination behavior.

The limited size of our recordings of 10-100 frequency-tuned neurons per animal precluded direct prediction of behavioral thresholds: an extrapolation from the measured population indicates that O(1000) neurons would be required to fully account for behaviorally observed frequency discrimination threshold (S2 Figure). This observation is consistent with an anatomical estimate suggesting that O(1000) neurons in the mouse AC are responsive to a given tone (mouse cortex has ∼10^5^ neurons/mm^3^ [37], the AC is ∼5 mm^3^ in size, ∼30-50% of neurons are frequency tuned, and the tuning width is ∼1/10 of the auditory spectrum). The fact that we were able to make strong predictions about optogenetic effects on behavior despite this sub-sampling suggests that the changes we observed in the measured cortical neurons were representative of changes occurring across the entire AC. Comprehensive recordings or imaging of more complete populations of AC neurons will enable a test of the hypothesis that frequency discrimination performance is limited by the encoding at the cortical level.

We observed a relationship between neural responses and behavior under the specific conditions of optogenetic manipulations of cortex, but the methods used here will be broadly useful for understanding other complex phenomena. For example, consider appetitive and aversive conditioning which have been shown to effect cortical remapping [31, 38, 39], while leading to diverse behavioral responses [9, 17]. In particular, such remapping leads to an overrepresentation of the aversive stimulus in cortex, but while some animals show improved frequency discrimination acuity, others are impaired. In fact, our results indeed show that different ways over-representing the aversive stimulus can lead to improved or impaired acuity, depending on the details of the change (Figure 6). For example, shifting the neural tuning curves towards the aversive frequency leads to increased sensitivity near this tone (Figure 6B), while simply increasing neural activity in response to tones near the aversive frequency can lead to impaired discrimination (Figure 6C). Fisher information analyses which we used in our paper have proven generally useful in understanding neural coding, providing insight about sound localization strategies [40], sensitivity to variations in sound levels [41], habituation to repeated sounds [42], detection of sound in noise [43] and heading perception [27]. Our method allows applications of this technique to compare behavioral discrimination thresholds to neurometric uncertainty in tone frequency estimation.

**Figure 6.**
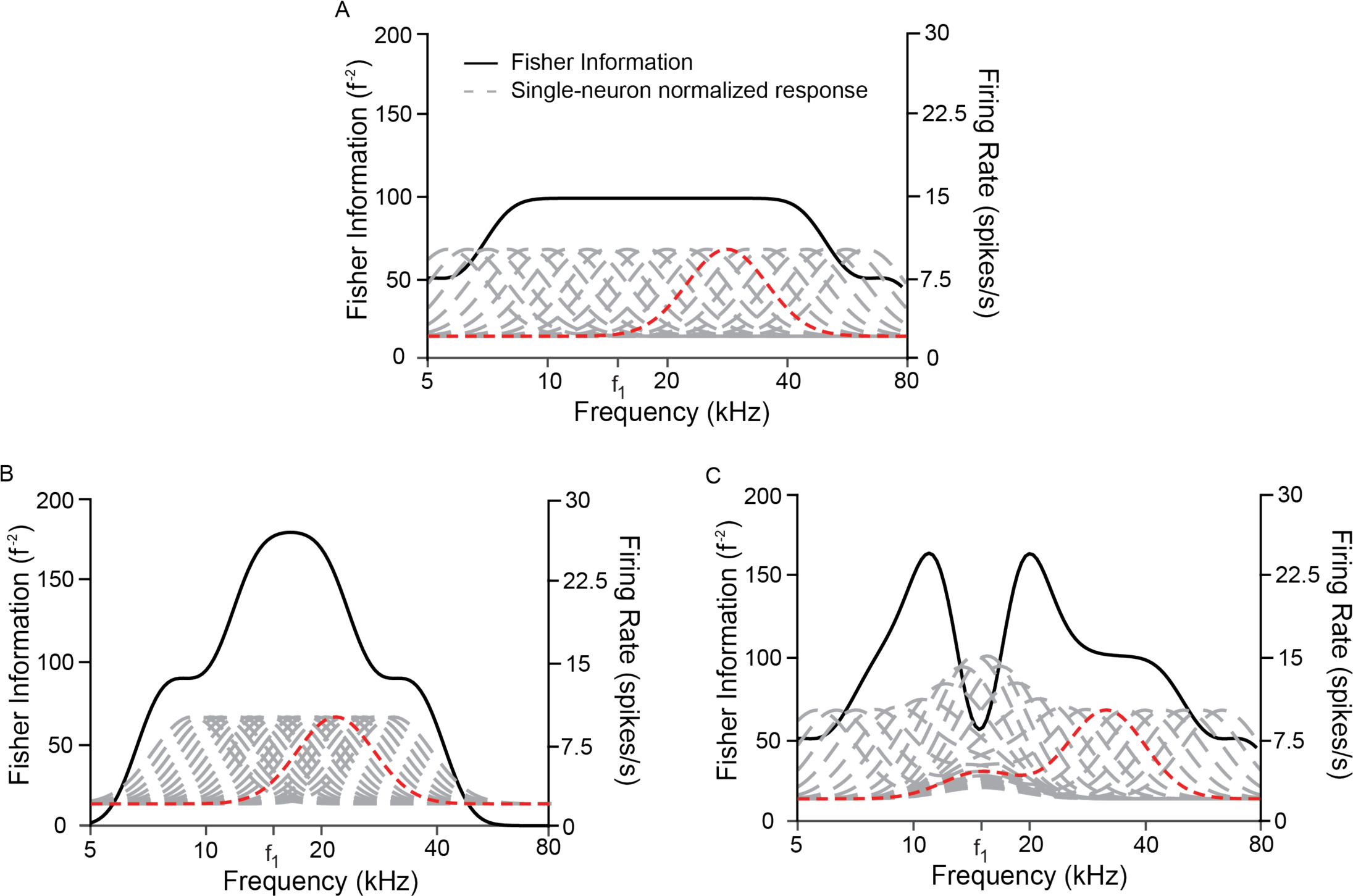
Overrepresenting a specific frequency can increase or reduce sensitivity to that frequency. **A**. Fisher information (black) computed from a homogeneous population of neurons (responses in gray) has an even sensitivity across a broad range of frequencies. A sample tuning curve (red) is used to illustrate neural transformations in **B** and **C**. Neurons have baseline activity of 2 spikes/s, peak response of 10 spikes/s, peak frequency spaced 1/20^th^ of a decade apart, with HWHM of .1 decades. **B**. Fisher information is plotted for a neural population overrepresenting frequency *f*_1_ by shifting peak frequencies halfway between their original location in **A** and *f*_1_. Fisher information approximately doubles near *f*_1_, but is reduced near the edges. **C**. Fisher information is plotted for a neural population overrepresenting frequency *f*_1_ by adding a Gaussian bump near *f*_1_ with an amplitude that diminishes with distance between the preferred frequency of the neuron and *f*_1_. Fisher information is *diminished* at *f*_1_, leading to reduced sensitivity at this frequency, despite its overrepresentation within the population firing activity.

Whereas we used responses to single tones as the stimulus in this study, AC neurons generally respond to more complex sounds in a manner that is not well explained by the single-tone responses [44, 45]. Our methods could be used to probe the fidelity of cortical representation and behavior in response to any parameterized space of auditory stimuli, e.g. auditory textures, phonemes, or overtone profiles. Similarly to work in visual texture perception [46, 47], one could test the discrimination thresholds along different dimensions of the texture space [48-50]. Changes in these thresholds due to optogenetic circuit manipulation could then be compared with changes in a neurometric threshold based on Fisher Information, as applied here. Such studies will elucidate the role of AC in facilitating complex auditory discrimination.

Viewed in aggregate in terms of a mean over individuals, our results would have yielded small average effects with large individual variations appearing as “noise’’. Averaging in this way would have obscured the function of AC in frequency discrimination because the correlation between individual circuit changes and individual behavioral changes would have been missed. It is only by treating the mice as individuals that we were able to understand the general role auditory cortex plays in shaping the response. We suggest that such attention to individual variation will be broadly important throughout neuroscience as large-scale recordings begin to reveal the neural basis of behavioral diversity.

## Methods

*Animals.* All experiments were performed in adult male mice (supplier - Jackson Laboratories; age, 12-15 weeks; weight, 22-32 g; PV-Cre mice, strain: B6; 129P2-Pvalbtm1(cre)Arbr/J; CamKIIa?-Cre: B6.Cg-Tg(CamKIIaα-Cre)T29-1Stl/J) housed at 28° C on a 12h light:dark cycle with water and food provided *ad libitum*, 5 or fewer animals per cage. All animal work was conducted according to the guidelines of University of Pennsylvania IACUC and the AALAC Guide on Animal Research. Anesthesia by isofluorane and euthanasia by carbon dioxide were used. All means were taken to minimize the pain or discomfort of the animals during and following the experiments. Experiments were performed as described previously [9].

*Viral constructs.* Modified AAV vectors were obtained from Penn VectorCore. To suppress PVs, we used modified AAV vector encoding Archaerhodopsin (Arch) under FLEX design (Addgene plasmid 22222, AAV-FLEX-Arch-GFP [51]. To activate either PVs iin PV-Cre mice and or excitatory neurons in CamKIIα-Cre mice, we used modified AAV encoding ChR2 under FLEX design (Addgene plasmid 18917 AAV-FLEX-ChR2- tdTomato, ChR2 [52]. As a control, we used modified AAV vectors encoding only GFP or tdTomato under FLEX design.

*Experimental methods overview.* Methods have been previously described [9]. Briefly, at least 10 days prior to the start of experiments, mice were injected with a viral construct, if any, and implanted with optical cannulas, and a headpost, as previously described, under isoflurane anesthesia. Viral construct injection targeted AC using stereotaxic map. Fiber-optic cannulas were implanted bilaterally over the injection site at depth of 0.5 mm from the scull surface, along the dorsal-ventral axis. After recovery, mice were habituated to the head-fixing apparatus, and subjected to behavioral frequency discrimination tests for 1-3 days, followed by electrophysiological recordings in the auditory cortex. On half of the trials in behavioral and neurophysiological recordings, light was presented through the fiber-optic cannula to activate or suppress target neurons. Upon conclusion of experiments, brains were extracted, fixed and subjected to immunostaining. Viral spread was confirmed postmortem by visualization of the fluorescent protein expression in fixed brain tissue, and its co-localization with PV or excitatory neurons, following immuno-histochemical processing with the appropriate antibody.

*Behavioral frequency discrimination.* Behavioral frequency discrimination was measured using a modified PPI procedure on daily basis [9, 53]. As previously reported [9, 17], PPI provides psychometric response curves for frequency discrimination over the course of a single session that lasts less than 1 h and does not require training the subject, which can confound interpretation of fear conditioning. Mice were head-fixed, connected to optical cannulas as needed, and placed on a load-bearing platform. Sound was presented through a speaker, consisting of a background tone (15kHz, 10-20 s, 80 dB SPL) that, on each trial, switched to a pre-pulse tone (10.2, 12.6, 13.8, 14.7, and 15.0 kHz, 60 ms, 80 dB SPL) followed by startle noise (SS, broad-band noise, 100 dB SPL, 20 ms). The frequency difference between the background and pre-pulse tone is denoted *Δf*. Each session started with presentation of 10 startle-only trials, to ensure that the mouse habituated to the beginning of the session. Data from these trials were not used for the analysis. Each pre-pulse tone was repeated in a pseudo-random order at least 5 times during each behavioral session.

The Acoustic Startle Response (ASR) for a given *Δf* was computed as the average over trials of the difference between the maximum vertical force applied within the 500 ms window following SS and the average baseline activity during 500 ms prior to SS. In each PPI session, the PPI was calculated as:

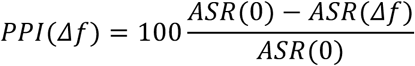

where *PPI* is reported in % relative to maximum *PPI*, and *ASR*(*Δf*) is measured using the 50% of the strongest ASR magnitudes for each PP frequency. 50% is chosen for consistency with previous work, as the psychometric curve has the steepest slope at this value [17]. Behavioral threshold was determined by fitting the PPI with a generalized logistic function, and defining the threshold for the fit as the *Δf* that produced 50% of the maximum PPI. Because the primary source of uncertainty related to how close the animal’s threshold was to sampled points, we computed thresholds for the set of fits producing less than a 25% increase in MSE relative to the best fit (although increasing this cutoff to 60% yielded small differences). We took the mean and standard deviation of the resulting set of thresholds to reflect the animal’s threshold. Each psychometric functions consisted of 5 data points representing the difference between the background frequency and 5 prepulse frequencies (PP). Each data point was obtained by averaging ASRs from all repetitions corresponding to a given frequency. In a standard PPI session, 20 repetitions of each PP were presented (100 trials in total). However, if either threshold was out of the range (0.5–32%) or the fit coefficient of the curve (R^2^) was below 0.7, the mouse underwent an additional 10 repetitions (50 trials). If threshold and fit curve failed to meet the above criteria after 200 trials, the session was excluded from statistical analysis (3 out of 61 sessions). Previous studies have shown that psychometric thresholds obtained from day-to-day measurements were stable [9, 17, 54]. Light-on trials included a 1s laser presentation that starts 0.5 s preceding PP onset. Light-off trials included laser presentation at quasi-random position during ITI. All analysis was performed separately on light-on and light-off trials. Simple randomization was used to assign the subjects to the experimental groups. A pseudorandom sequence was used for tone presentation during PPI tests. During PPI procedure, the timing of the laser presentation on Laser-off trials was pseudo-randomized with respect to the timing of the tones. Blinding of experiment with respect to animal groups was not possible as animals in different groups underwent different experimental protocols.

*Neuronal tone response measurement.* All recordings were carried out inside a double-walled acoustic isolation booth (Industrial Acoustics) as previously described [9]. Activity of neurons in the primary auditory cortex of head-fixed, awake mice was recorded via a silicon multi-channel probe (Neuronexus). Putative principal (excitatory) neurons were identified using waveform and spontaneous firing rate (for details see [9]). Acoustic stimulus was delivered via a calibrated magnetic speaker (Tucker-Davis Technologies) [45]. We measured the frequency tuning curves by presented a train of 50 pure tones (50ms long, ISI 450 ms) with frequencies spaced logarithmically between 1 and 80 kHz and at 8 intensities (sound pressure levels, SPLs) spaced uniformly between 10 and 80 dB, a standard procedure in characterizing auditory responses, and determined the threshold amplitude for tone at each frequency for each neuron [9, 45]. For data analysis we averaged responses of the neurons to each tone across 3 highest amplitudes. Each tone was repeated twice in pseudo-random sequence and the stimulus was counter-balanced for laser presentation. On light-on trials, light was presented via the optic cannulas, with the onset of 100 ms prior to tone onset, and lasting for 250 ms. The full stimulus was repeated 5 times.

*Histology.* Virus spread was confirmed postmortem by visualization of the fluorescent protein expression in fixed brain tissue, and its colocalization with PV or excitatory neurons, following immuno-histochemical processing with the appropriate antibody.

*Identification of Putative Excitatory Neurons.* Putative principal (excitatory) neurons were identified using waveform and spontaneous firing rate (for details see Aizenberg et al., 2015).

*Neural Response Analysis*. The neural frequency response function was calculated using the average frequency tuning curve across the three highest intensities. The results were modeled using a Gaussian in log-frequency space:

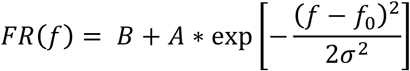

where *B* represents the baseline response, *A* represents the amplitude of the strongest evoked response (relative to baseline), *f*_0_ represents the frequency evoking the strongest response, and *σ* corresponds to the width of the frequency response function. Only neurons whose Gaussian fit had *R*^2^ > .6 were kept for further analysis.

In order to calculate Fano factor of a neuron, we calculated mean and variance of the firing rate to each combination of frequency and SPL. The slope of these data was taken as an estimate of Fano factor.

The correlation between neurons (calculated only between simultaneously recorded neurons) was computed using a reduced measure of deviation from the mean for each neuron:

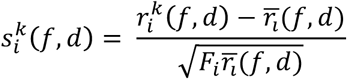

where *k* denotes the repetition number (1-5), *i* denotes the neuron, *f* denotes the frequency, *d* denotes the intensity, *r* denotes the evoked response, 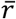 denotes the average response (firing rate) of a neuron to a particular frequency and intensity, and *F* denotes the Fano factor of that neuron. For a generalized Poisson process, *s_i_* has zero mean and unit variance because the variance is proportional to the mean. The correlation between two neurons is then given by

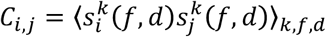

*Computing Fisher Information.* Fisher information was calculated to provide a measure of neurometric frequency discrimination. Fisher information was calculated numerically from the recorded data based on the characterization of the neural responses (see *Neural Response Analysis*), and is given by:

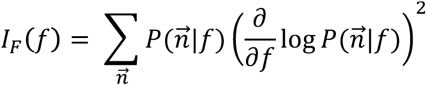

where *P* is the probability that the population of neurons produces 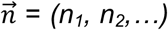 spikes in response to the tone *f*. Assuming Poisson variability and independent neurons,

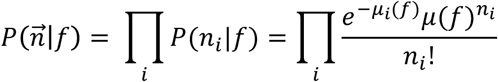

where the first equation expresses the independent neuron assumption and the second step uses the assumption of Poisson variability. Here, *μ_i_* and *n_i_* are the expected and observed number of spikes from neuron *i*, respectively. For mice where multiple recording sessions were performed, neurons were pooled across sessions. A second model assumes independent neurons and uses a generalized Poisson distribution:

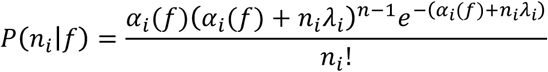

where 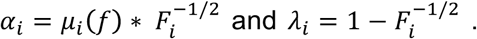 This model only allows for Fano factors above 1, which is consistent with other models of neural variability and measured cortical variability [24, 26]. We therefore set any measured Fano factor less than 1 to exactly 1 for the purpose of the Fisher information calculation.

*Estimating the number of neurons for neurometric thresholds.* Animals displayed varying levels of frequency discrimination acuity, and we measured different sized subsets of frequency-tuned neurons in each experiment. In order to estimate the number of neurons required to account for behavioral discrimination acuity from the measured population, we utilized the fact that Fisher information for a population of independent neurons is the sum of the Fisher information from each neuron.

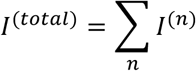

We then assume that the population of neurons that we measured are representative of the overall population (at least the population that contributes to discrimination at the relevant frequency).

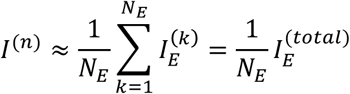

Here we use *n* to refer to the overall population that we seek to estimate, *k* to index the neurons that were measured, *N_E_* is the number of frequency-tuned neurons measured in a specific mouse, 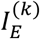 is the Fisher information of these experimentally measured neurons, and 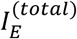 is the total information used to calculate the neurometric threshold. Recalling that the neurometric threshold is defined as *I*^−1/2^, we then have

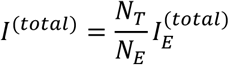

We can rewrite this as

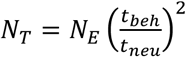

where *t_beh_* is this behaviorally measured threshold and *t_neu_* is the neurometric estimate of the threshold. This is plotted for each animal used in our analyses in S2 Figure, and we find that the average for both light-on and light-off conditions is just about 10^3^ neurons.

*Comparing behavioral and neurometric thresholds.* Neural responses were accumulated over all recording sessions for each mouse, and used to compute the Fisher information for the animal. The Fisher information provides a bound on the variability of an unbiased estimator, and any criterion level of decoding performance scales with the inverse-square root of the Fisher information. If neural responses are independent, the Fisher information scales linearly with the number of neurons. In order to compare threshold predictions between mice with different numbers of measured neurons, we computed the average Fisher information per neuron. This allowed us to compare across mice and to estimate the minimum number of effective frequency-tuned neural units that must contribute to explain the observed frequency discrimination performance [18]. To estimate the neurometric threshold, we first computed the inverse square root of the Fisher information per neuron, and took an average over a small region around 15 kHz (the baseline frequency used in behavioral tests). Only mice with more than 10 recorded frequency-tuned neurons were included in the analysis (19 mice). Our estimate is a lower limit on the uncertainty in the estimate of this tone frequency based on the recorded population responses. Since Fisher information scales linearly in an independent population, this method also provides an estimate on the effective number of neurons which must contribute to the tone representation in order to account for behavioral discriminability (S2 Figure). The observed neurometric discriminability with order one thousand independent neurons is similar to behavioral discrimination acuity.

*Data availability.* All relevant data will be deposited in the Dryad database and made publically available.

*Code availability.* All relevant code will be posted on Github and made publically available.

## Supplementary Information

**S1 Table:**
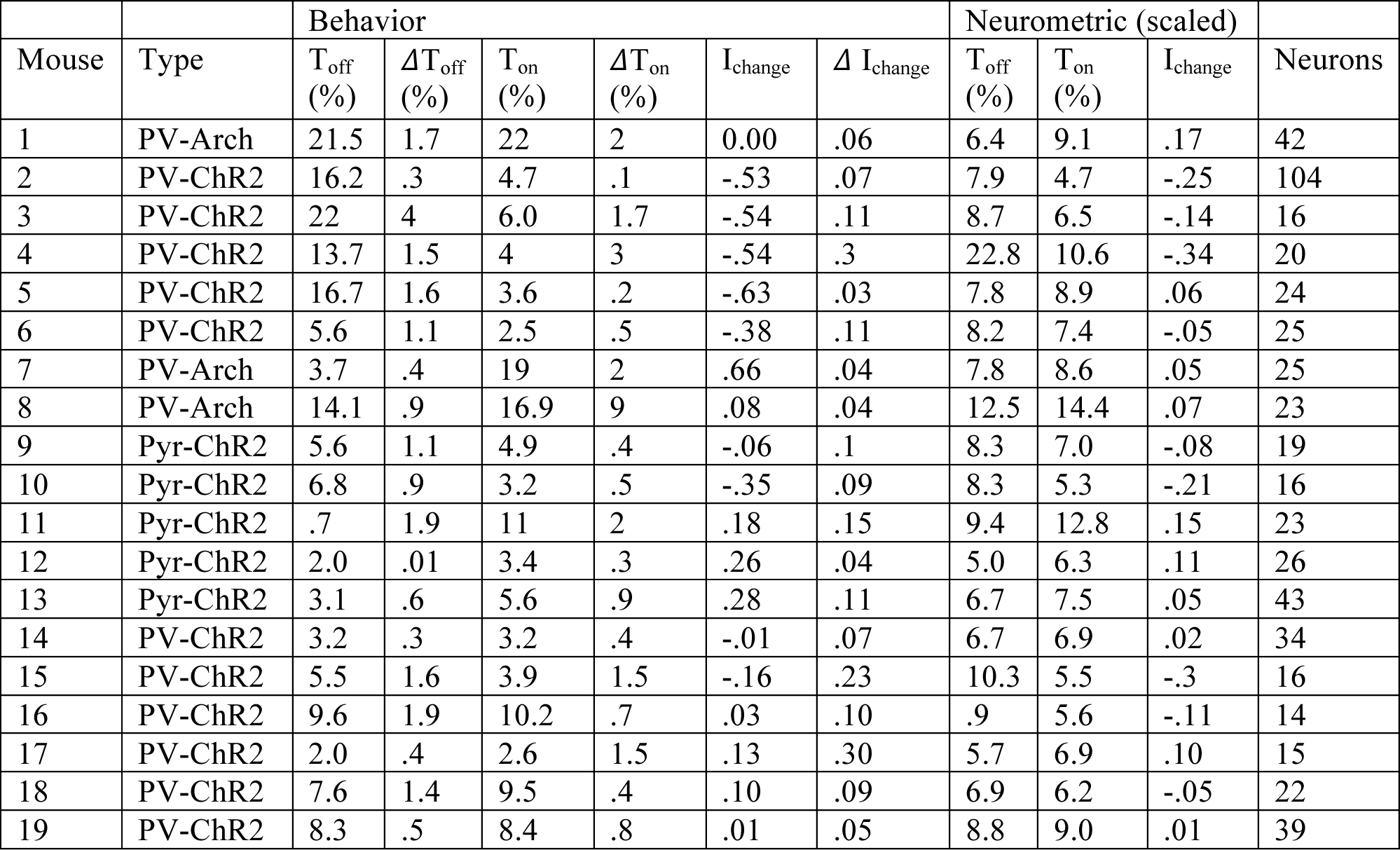
Table containing mouse identity, type, relevant behavioral and neurometric thresholds, as well as number of recorded neurons. Neurometric threshold computed using Fisher information is scaled to an effective population size of 1000 neurons to control for differences in numbers of measured neurons.

**S1 Figure:**
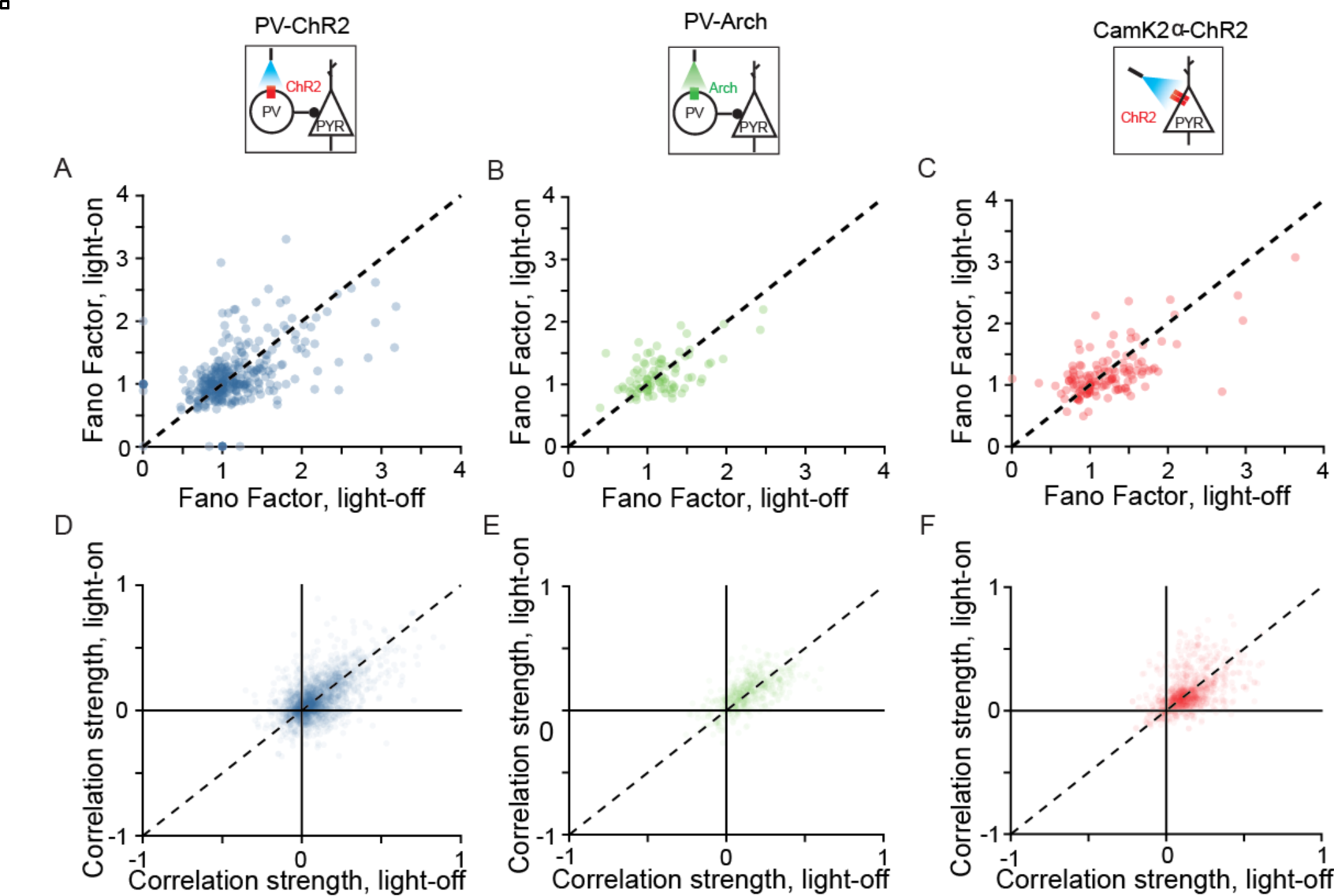
Fano factor and correlation scatter plots comparing light-on and light-off conditions. **A**-**C**: Fano factor with and without light on for PV-ChR2, PV-Arch, and Pyr-ChR2 mice, respectively. **D**-F: Pairwise correlations with and without light on for PV-ChR2, PV-Arch, and Pyr-ChR2 mice, respectively.

**S2 Figure.**
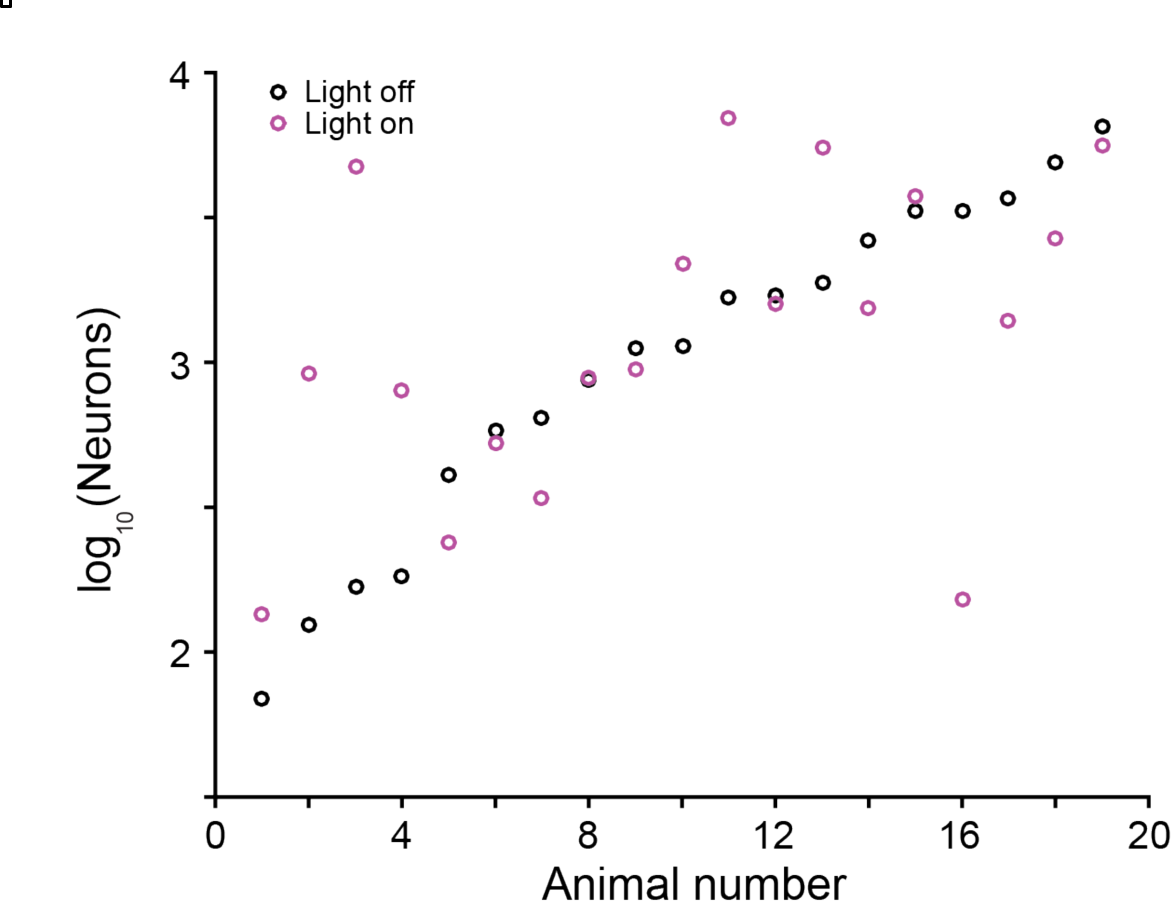
Number of frequency-tuned neurons required to account for behavioral sensitivity for each mouse. Average of both light-on and light-off conditions is ∼10^3^ neurons.

